# Early life stress impairs social function through AVP-dependent mechanisms

**DOI:** 10.1101/741702

**Authors:** Nichola M. Brydges, Jessica Hall, Caroline Best, Lowenna Rule, Holly Watkin, Amanda J. Drake, Catrin Lewis, Kerrie L. Thomas, Jeremy Hall

**Affiliations:** Neuroscience and Mental Health Research Institute, Cardiff University, Hadyn Ellis Building, Maindy Road, Cardiff, CF24 4HQ, UK; National Centre for Mental Health, Cardiff University, Hadyn Ellis Building, Maindy Road, Cardiff, CF24 4HQ, UK; University/BHF Centre for Cardiovascular Science, University of Edinburgh, Edinburgh, EH16 4TJ UK; School of Biosciences, Cardiff University, Museum Avenue, Cardiff, CF10 3AX, UK; MRC Centre for Neuropsychiatric Genetics and Genomics, Cardiff University, Hadyn Ellis Building, Maindy Road, Cardiff, CF24 4HQ, UK

**Keywords:** Early life stress, social behaviour, arginine vasopressin, oxytocin

## Abstract

Impaired social function is a core feature of many psychiatric illnesses. Adverse experiences during childhood increase risk for mental illness, however it is currently unclear whether stress early in life plays a direct role in the development of social difficulties. Using an animal model of pre-pubertal stress (PPS), we investigated effects on social behaviour, oxytocin and arginine vasopressin (AVP). We also explored social performance and AVP expression in participants with borderline personality disorder (BPD) who experienced a high incidence of childhood stress. Social behaviour was impaired and AVP expression increased in animals experiencing PPS and participants with BPD. Behavioural deficits in animals were rescued through administration of the AVP receptor 1a antagonist Relcovaptan (SR49059). AVP levels and recognition of negative emotions were significantly correlated in BPD participants only. In conclusion, early life stress plays a profound role in the precipitation of social dysfunction, and AVP mediates at least part of this effect.

## Introduction

Altered social function is a core component of several adult psychiatric illnesses. For example, depression and schizophrenia are commonly associated with social withdrawal, while major personality disorders can be associated with unstable social relationships (borderline personality disorder, BPD) or aggressive interactions (antisocial personality disorder)^1-4^. Impairments in social behaviour can also be a key determinant of functional outcome in these conditions^5-7^. However, relatively little is known about the causes of impaired social function in psychiatric conditions and their relationship to important aetiological factors associated with psychiatric disorders such as developmental stressors.

Early life stress has been shown to impact on social behaviour and functioning in both animal and human studies ^8-13^. In rodent models stress at different developmental time points is associated with altered social interactions in adulthood, with the exact nature and timing of the stressor often influencing later outcomes^12^. The majority of studies have focussed on stressors early in development such as maternal separation, but later stressors also exert an effect, with non-social peri-pubertal stress resulting in aggression in adulthood, and physical pre-pubertal stressors impacting on later social interaction ^12, 14-21^. In humans, adverse early life experiences have been strongly associated with a range of later difficulties in social interaction in longitudinal and cross-sectional studies^8, 9, 22, 23^. In particular, childhood and adolescent stressors have been linked to later social anxiety, withdrawal and aggressiveness, although there is substantial variation in outcomes and not all exposed individuals are affected ^24-28^. There is also substantive evidence that childhood adversity is associated with an increased risk for psychiatric disorders in which altered social interaction is a prominent feature including social anxiety, depression and personality disorders^29-31^. A particularly strong association between developmental adversity and later illness is seen in the case of BPD in which impaired social interactions are a core feature of the presentation^30, 32-36^.

The mechanisms through which early life stressors impact later social behaviour are still largely unknown. The evolutionarily conserved social neuropeptides oxytocin (OXT) and arginine vasopressin (AVP) are known to play an important role in social behaviour across species ^37-41^. AVP and OXT are evolutionarily ancient nonapeptides found in various guises throughout the animal kingdom ^42^. In mammals, they are predominantly manufactured in the paraventricular and supraoptic nuclei of the hypothalamus, and released into peripheral circulation via the pituitary gland ^37^. OXT and AVP play a wide variety of roles in social behaviour in animals, ranging from inter-male and maternal aggression, conspecific affiliation, social cognition and sexual behaviour^37,39,40^. In humans, intranasal administration of both OXT and AVP influence emotion processing and social cognition, and genetic and functional neuroimaging studies demonstrate a role for OXT and AVP on social behaviour and related limbic brain circuitry^43, 44^.

Maternal separation in rodents has been shown to increase AVP levels specifically in the paraventricular nucleus and alter OXT levels and OXT/AVP receptor binding in an age and sex specific manner in the offspring^14, 45-47^. However less is known about the effects of stressors applied pre-pubertally, a period which may be more analogous to human childhood, on social peptide regulation. In humans, experience of childhood maltreatment has been associated with decreased levels of OXT in women and men, but the effects of early stress on AVP levels are less well known^48^.

Here we report three experiments investigating the relationship between early life stress, AVP/OXT and social behaviour. The aim of experiment 1 was to investigate the effects of pre-pubertal stress on subsequent social behaviour and levels of OXT and AVP in adult animals. Experiment 2 sought to determine whether changes in social behaviour resulting from PPS could be reversed thorough AVP receptor antagonism, using the AVP receptor 1a (AVPR1a) antagonist Relcovaptan (SR 49059). The third and final experiment assessed peripheral levels of AVP in patients with BPD (a group with a high incidence of childhood adversity) and controls and related these to social behaviour.

## Materials

### Experiments 1 & 2

#### Subjects

Female and male Lister Hooded rats were bred in house from adult pairs (Charles River) at Cardiff University. Litters were weaned on postnatal day (PND) 21 and housed in same sex cages (32cm × 50cm × 21cm) with littermates. Cages were lined with wood shavings, a cardboard tube and wooden stick were provided as enrichment, light was maintained on a 12:12-h light/dark cycle and food and water were provided *ad libitum*. All animal experimental methods were carried out in accordance with relevant guidelines and regulations of the European regulations on animal experimentation (Directive 2010/63/EU) and the UK Home Office Animals (Scientific Procedures) Act 1986. All experimental protocols were approved by the local ethical review body (AWERB) of Cardiff University. Eighty rats (male: 20 control, 20 PPS; female: 20 control, 20 PPS) were used for social testing in experiment 1, and one hundred and forty four (female: 22 control & vehicle, 24 control & relcovaptan, 20 PPS & vehicle, 30 PPS & relcovaptan; male: 12 control & vehicle, 14 control & relcovaptan, 10 PPS & vehicle, 12 PPS & relcovaptan) in experiment 2. In a separate cohort of animals (not subjected to behavioural testing), forty rats (male: 12 control, 10 PPS; female: 8 control, 10 PPS) were used for plasma collection (peripheral AVP), and a further 44 rats (male: 10 control, 13 PPS; female: 10 control, 11 PPS) for immunohistochemistry (central AVP).

#### Pre-pubertal stress

Pre-pubertal stress (PPS) was given to half of the litters on PND 25-27. This protocol has been described previously^49-51^, and was originally described by Jacobson-Pick and Richter-Levin^52^. Briefly, animals were given a 10 min swim stress in an opaque swimming tank (25cm high, 34cm diameter), 12 L capacity filled with 6L of 25±1°C water on PND 25, three sessions of 30 minute restraint stress (separated by 30 min breaks in the home cage) in plastic restraint tubes (15cm length, 5cm diameter) on PND 26, and three 30 minute elevated platform exposures (separated by 60 min breaks in the home cage) on elevated platforms (15×15cm, 115cm high) on PND 27. Stressors took place in a designated room, separate from the holding room. After PPS, animals were returned to their home cages and holding rooms and left undisturbed (aside from cage cleaning) until early adulthood (PND 60-67). Litters were randomly allocated to experimental groups (PPS or control). A minimum of 5 litters per group were used to minimise the effects of pseudo-replication, and litter of origin was accounted for in all statistical analyses.

#### AVPR1a antagonist

Relcovaptan (AVPR1a antagonist SR49059, Axon Medchem BV, The Netherlands) was dissolved in 15% dimethyl sulfoxide (DMSO) and 2% Tween 80 in 0.9% saline and administered intraperitoneally at a volume of 2mL/kg. The dose selected was 1mg/kg, as this dose reliably inhibits prosocial and autonomic effects of peripherally administered AVP^53, 54^. Vehicle was 15% DMSO and 2% Tween 80 in 0.9% saline.

#### Social testing

Testing took place in a clear acrylic tank (65cm×65cm×40cm high) placed on the floor in the centre of a dimly lit room (30 lux). Interactions were filmed from above, and a microphone was suspended above the tank, connected to Avisoft SASLab Pro (avisoft bioacoustics, Germany) to capture ultrasonic vocalisations, which are frequently emitted during murine social encounters. Videos were then watched back by an observer blind to group, and a range of parameters were recorded, including latency to initial contact, duration of each contact, number of contacts and total contact time. Recorded vocalisations were analysed with Avisoft SASLab Pro program. Very few 22 kHz vocalisations were produced, so all analyses are based on 50 kHz vocalisations. In rats, 22 kHz vocalisations are most often emitted in aversive situations (e.g. presence of predators, pain), whereas 50 kHz reflect more positive affective states (e.g. during social contact, mating, in response to drugs of abuse or food)^55^.

Three hours before testing, animals were single housed in their holding room, to increase the desire for social contact. One hour before testing, they were transferred to the testing room, to habituate them to this environment. In experiment 2 only, half of the animals from each group were given 1mg/kg Relcovaptan via intraperitoneal injection 30 minutes before testing, the remaining half were administered vehicle only. During testing, two stranger animals were placed into opposite sides of the arena, facing the wall, and allowed to freely interact for 15 minutes. After this time they were returned to their home cages. The arena was cleaned with ethanol wipes between pairs of animals. Animals were tested in same-sex pairs, with each member of a given pair originating from the same group and treatment condition (PPS or control, vehicle or Relcovaptan) but different litters, so the animals had not previously met. Each pair was treated as one experimental unit for behavioural analysis.

#### Tissue & plasma collection

*Experiment 1*. For plasma AVP and OXT analyses, animals were sacrificed at PND 60 using a rising concentration of CO_2_, decapitated and trunk blood was collected using EDTA microvette collection tubes (Sarstedt, Germany). Blood was spun at 1500 × g for 10 minutes, plasma was removed and stored at −20°C. For immunohistochemistry, animals were killed at PND 60 by transcardial perfusion with 0.01M PBS and 4% paraformadelhyde (PFA) under anaesthesia for immunohistochemical analysis of AVP in the supraoptic and paraventricular nuclei. Brains were left in PFA overnight (4°C), transferred to 30% sucrose solution for cryoprotection. Coronal 30µm sections were cut through the entire hypothalamic extent on a freezing microtome (Leica RM2245) and placed into a solution of cryoprotectant for storage at −20°C until immunohistochemical analysis. *Experiment 2*. After social testing animals were culled (half of the animals were culled 20 minutes after testing (direct), the rest one week later (delayed)) and trunk blood samples taken as in *experiment 1*.

#### ELISA assays

For experiment 1, rat plasma samples were assayed untreated. For experiment 2 and all human samples, plasma samples were dried down by adding 2:1 ice cold acetone:plasma, centrifuging at 3000xg for 20 minutes, transferring the supernatant, adding 5x supernatant volume of ice cold petroleum ether to the supernatant, centrifuging at 3,000xg for 10 minutes, discarding the top ether layer and drying the remaining aqueous layer under nitrogen gas. This remaining pellet was reconstituted with 250ul of assay buffer and used for ELISA analysis. Arg^8^-vasopressin and OXT (Enzo Life Sciences, UK) ELISAs were conducted according to the manufacturer’s instructions.

#### Immunohistochemistry

Three sections per animal per region (supraoptic nucleus, paraventricular nucleus) were stained for AVP. These were matched for bregma between different animals. Sections were washed between each step for 3 × 5 minutes in 0.01M Tris-buffered saline (TBS, pH 7.4) and all steps were carried out at room temperature unless otherwise specified. Sections were blocked with BLOXALL (Vector laboratories, UK) for 10 minutes, then blocking solution (2% goat serum, 0.3% Triton-X in 0.01M TBS) for 60 minutes, rabbit anti-AVP (1:7500 in blocking solution, Millipore, UK) for 24 hours followed by biotinylated goat-anti-rabbit (3ug/ml in blocking solution) for 45 minutes, then 30 minutes in VECTASTAIN ABC reagent (avidin-biotinylated horseradish peroxidase complex, Vector Laboratories, UK), before developing in diaminobenzidine solution (Vector Laboratories, UK) for 5 minutes. Washed sections were mounted onto glass microscope slides and coverslipped with Vectamount (Vector Laboratories, UK). Slides were then imaged at 20x using Axio Scan Z1 (Zeiss). The optical density of AVP immunoreactive cells in the supraoptic and paraventricular nuclei were quantified as grey density per area minus background in digitized images using Zen Blue software (Zeiss).

#### Data analysis

Data were analysed using generalised linear models in JMP (statistical software, SAS Institute, Cary, NC, USA). Group (control or PPS), sex, Relcovaptan/vehicle (experiment 2 only), direct or delayed sacrifice (experiment 2 only) and all interactions were fitted as factors, and latency to contact, total contact time, average contact duration, number and duration of vocalisations, AVP and OXT were fitted as responses.

### Experiment 3

#### Participants

A total of 20 people with borderline personality disorder (BPD) were recruited from the National Centre for Mental Health (NCMH, Welsh Government funded research centre) participant pool database. Patients with BPD were chosen as this condition is typically associated with very high levels of childhood trauma^56^. A total of 18 healthy age and sex and matched controls with no psychiatric disorder were recruited from the community. Exclusion criteria for all participants included current substance dependence, neurological illness, psychotic disorder diagnosis (bipolar I or schizophrenia) and pregnancy, and additionally no psychiatric disorder for control participants. The BPD group consisted of 16 females and 4 males, mean age 43.4 (range 25-71), controls 14 females and 4 males, mean age 38.4 (range 20-64). In the BPD group, 11 were being treated with antipsychotic medication and 13 were being treated with antidepressant medication. BPD diagnosis was confirmed in the BPD group and excluded in the controls using the SCID-II (Structured Clinical Interview for DSM Disorders II). Current symptoms of depression were rated using the Hamilton Rating Scale for Depression (HADS). Participants also completed the Childhood Trauma Questionnaire (CTQ), a self-report measure of incidence and severity of childhood trauma consisting of 28 statements relating to one of five subscales of neglect (physical and emotional) or abuse (emotional, physical and sexual). The study was approved by the NHS research ethics committee, and all participants gave informed, written consent, had the opportunity to discuss the study and understood they were free to withdraw at any point.

#### Social testing

Participants were given the Ekman 60 faces task, which assess overall emotion recognition performance and identification of basic emotions. During testing, 60 different faces were presented one by one on a computer screen for 3 seconds, depicting the faces of actors portraying one of six basic emotions (fear, disgust, anger, sadness, happiness and surprise). Ten incidents of each emotion were portrayed during the task. Participants were given as long as needed to identify the emotion. The response choices were shown on the screen, and participants made their selection using a computer mouse. Responses were recorded automatically. Faces were selected from the Ekman and Friesen series of Pictures of Facial Affect^57^, a widely used and validated series of photographs in facial expression research.

#### Sample collection and analysis

Participants were visited in their own homes. Blood samples were collected from participants before questionnaire administration and social testing. In total, blood samples were obtained from 17 BPD and 15 controls. Samples were placed in cool bags containing reusable freezer blocks and transferred to the laboratory where they were centrifuged at 16000xg for 15 minutes at 4°C. The cell-free supernatant plasma fraction was removed and transferred to −80°C storage until analysis. AVP was analysed in plasma following the same protocol as for animal samples, using human Arg^8^-vasopressin ELISA kit (Enzo Life Sciences, UK). Copeptin was analysed using human copeptin ELISA (Novus Biologicals, UK), according to manufacturers instructions.

#### Data analysis

Data were analysed using generalised linear models or Mann-Whitney U tests in JMP (statistical software, SAS Institute, Cary, NC, USA). Data was checked for normality and homogeneity of variance and transformed to fit these assumptions when necessary. If data could not be transformed, non-parametric tests were used. For AVP and copeptin, group was fitted as factor with AVP/copeptin as a response. For the Ekman task, group (control or BPD), emotion (fear, disgust, anger, sadness, happiness and surprise) and group*emotion were fitted as factors, and number of correct responses as the response. Subject was nested within group and fitted as a random factor to account for repeated measures. Due to non-normal, non-homogenous data, Mann-Whitney U tests were used to analyse responses to the HADS and CTQ questionnaires. Spearman’s Rho was used to assess correlations between behavioural and AVP measures.

## Results

### Experiment 1

In experiment 1, we investigated the effects of pre-pubertal stress (PPS) on social behaviour and AVP expression in adult rats. PPS decreased latency to initial social contact (F_1,35_=4.27, p=0.046), decreased duration of each individual contact bout (F_1,35_=7.68, p=0.01), and decreased number (F_1,36_=10.17, p=0.029) and duration (F_1,36_=7.78, p=0.008) of positively valanced ultrasonic vocalisations (50kHz), and shortened maximum duration of vocalisations (F_1,36_=4.13, p=0.049), (Figure 1a-d) in males and females. There was no effect on the frequency of contacts made (F_3,35_=0.77, p=0.52).

**Figure 1.**
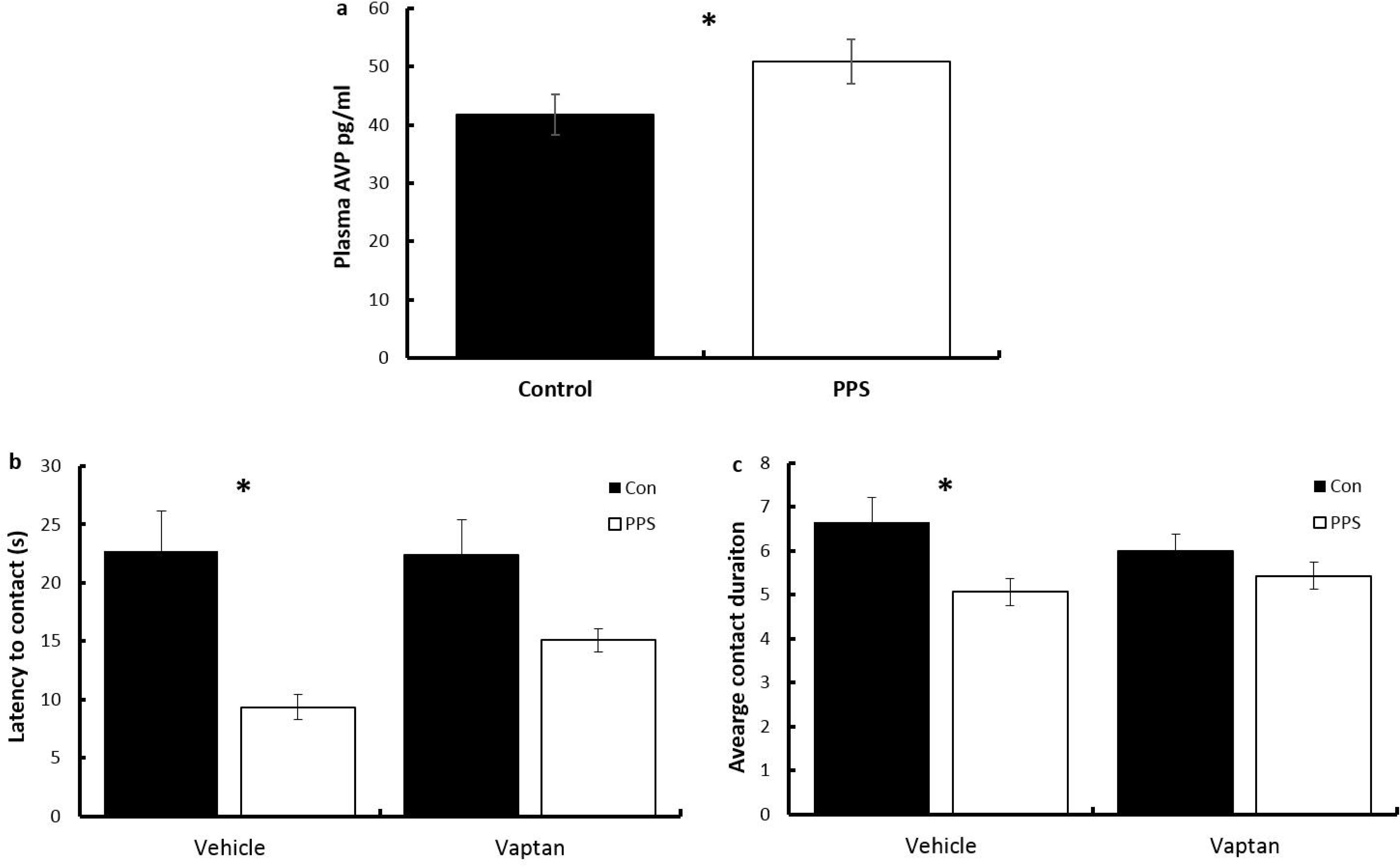
Social behaviour after PPS. PPS animals were a) faster to initiate contact, b) displayed shorter duration of contacts, c) emitted fewer vocalisations which were d) of shorter duration. *p<0.05, **p<0.01. Con=control, PPS=pre-pubertal stress. Error bars represent 1 S.E.

In separate cohorts (not behaviourally tested) PPS resulted in higher AVP levels in plasma (F_3,36_=2.07, p=0.04, Figure 2a) and centrally in the supraoptic (F_1,39_=5.31, p=0.03, Figure 2b, 2d) but not paraventricular (F_1,38_=0.23, P=0.63, Figure 2c, 2e) nucleus of the hypothalamus as assessed by immunocytochemistry. PPS did not influence plasma levels of OXT (F_3,36_=0.01, p=0.9).

**Figure 2.**
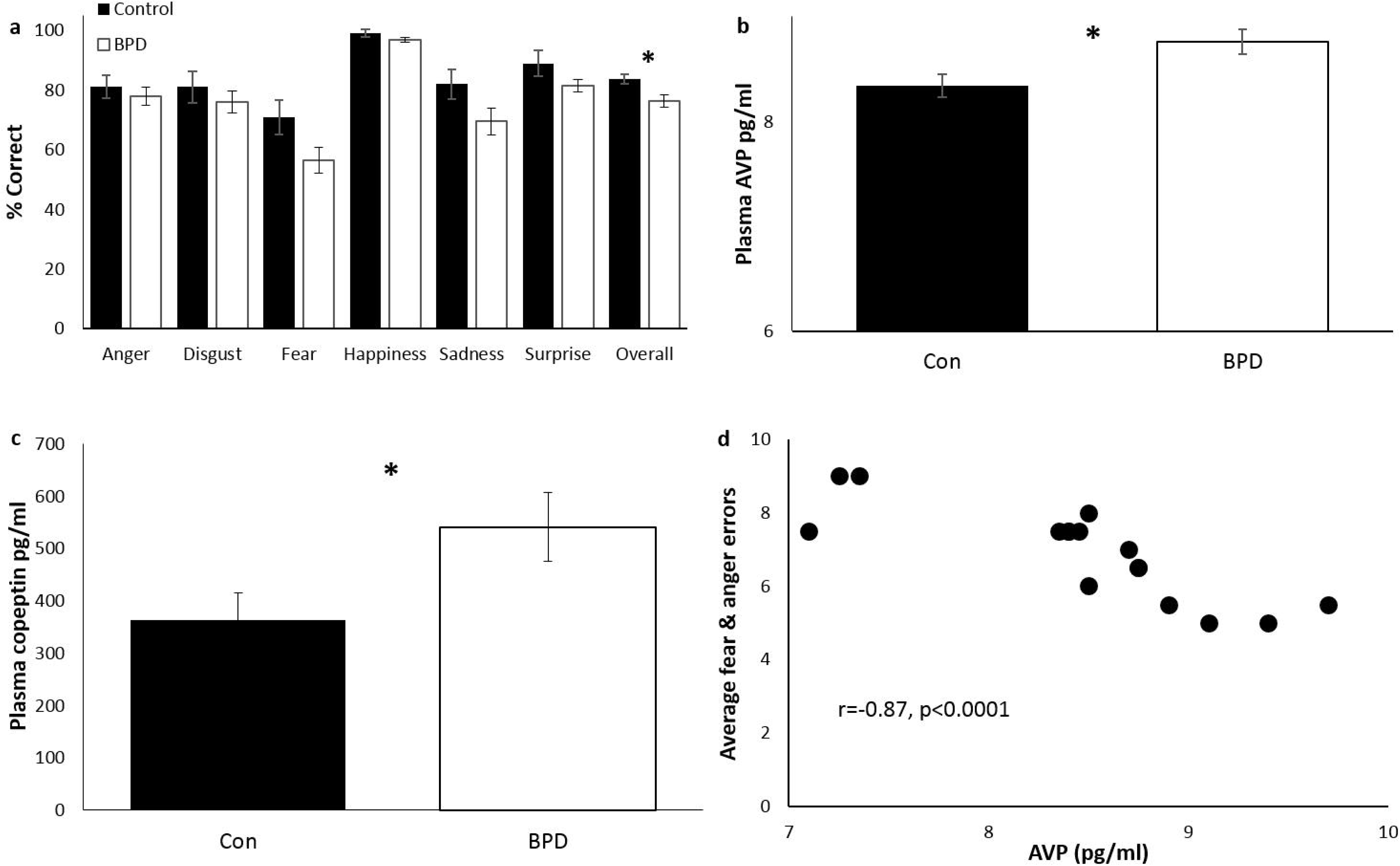
AVP levels after PPS. Following PPS, levels of AVP were elevated in a) plasma and b) supraoptic nucleus but not c) paraventricular nucleus. *p<0.05. Error bars represent 1 S.E.

### Experiment 2

In experiment 2, we investigated the ability of an AVPR1a antagonist, Relcovaptan, to rescue behavioural changes induced through PPS in our animal model. As in experiment 1, PPS significantly increased plasma levels of AVP in these animals (F_1,14.46_=6.95, p=0.01 Figure 3a). Replicating our findings in experiment 1, PPS reduced latency to initial contact and decreased duration of each individual contact during a social test. Administration of Relcovaptan substantially reduced this effect (latency: F_3,64_=3.57, p=0.02, contact: F_3,64_=3.1, p=0.03, data box-cox transformed, Figure 3b,c).

**Figure 3.**
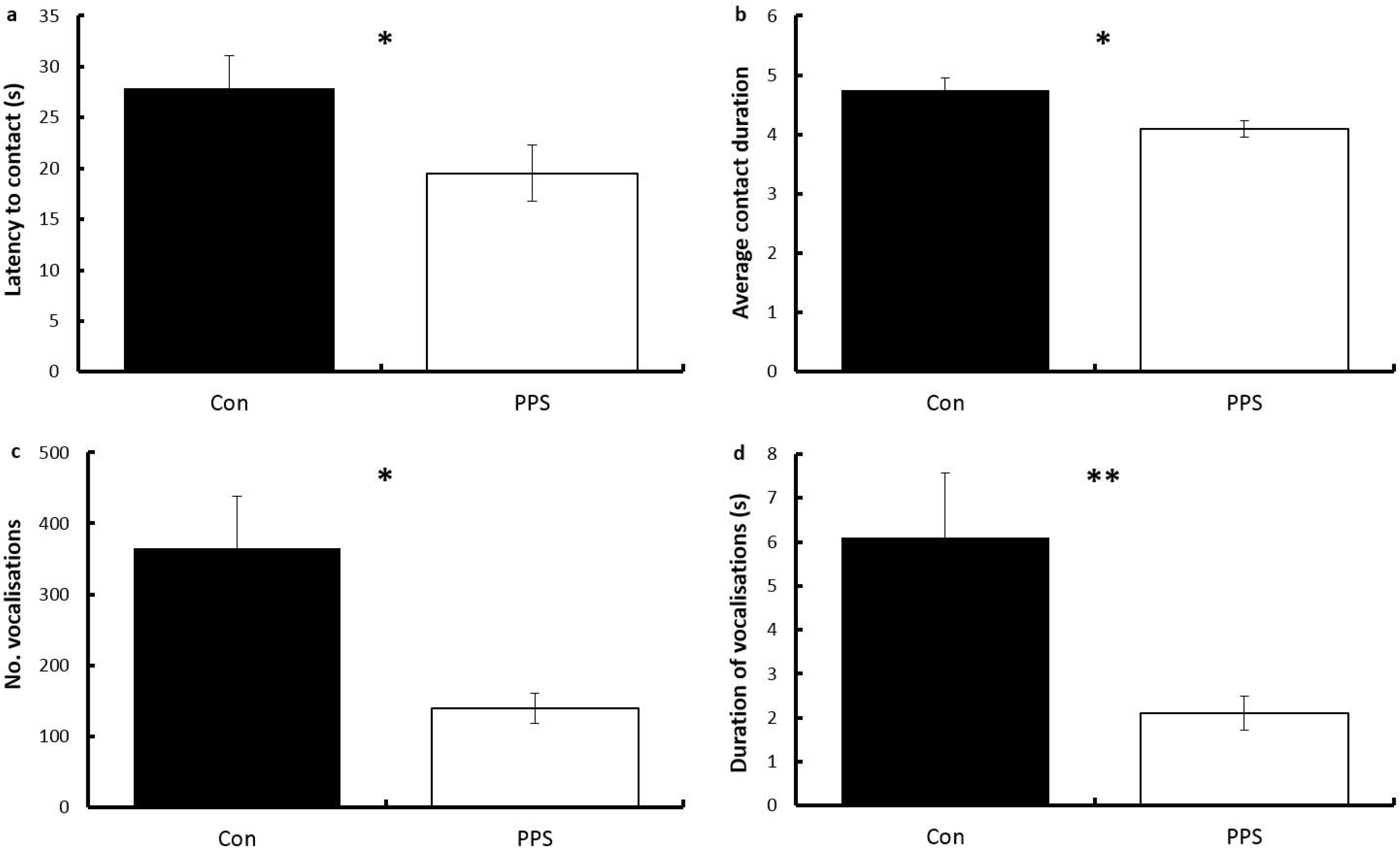
Effects of Relcovaptan on social behaviour after PPS. a) Following PPS plasma levels of AVP were again elevated. PPS animals were b) faster to initiate contact, this was reversed by administration of Relcovaptan, and c) displayed shorter duration of contacts, which was again reversed by Relcovaptan. *p<0.05. Error bars represent 1 S.E.

### Experiment 3

We next investigated AVP levels and social cognition (assessed using the EKMAN 60 faces task) in participants with BPD, a condition strongly associated with early life stress. There was no significant difference in age or sex between control and BPD groups (F_1,361.1,_ p=0.3). Participants with BPD scored higher on the Hospital Anxiety and Depression Scale (HADS) (S=192, p<0.0001) and Childhood Trauma Questionnaire (CTQ) (F_1,36_=45.34, p<0.001) across all physical and emotional abuse domains. Overall, BPD participants were worse at recognising emotion in the EKMAN task (F_1,36_=7.26, p=0.01, Figure 4a), replicating some previous findings but contrasting others ^58-60^. This was not specific to any emotion, as demonstrated by the lack of group*emotion interaction (F_1,180_=0.83, p=0.53). Plasma levels of AVP were higher in BPD compared to control participants (F_1,28_=6.42, p=0.02, Figure 4b), reflecting the higher peripheral AVP levels seen in the rat PPS model. Measurement of AVP has been suggested as problematic due to rapid clearance from circulation, instability in plasma and high platelet binding^61^. Therefore we also measured plasma levels of copeptin, the c-terminal segment of the AVP precursor peptide, which has been suggested a more stable, surrogate marker of AVP release^62^. Copeptin was also elevated in the BPD patients (F_1,24_=4.36, p<0.05, Figure 4c). After correction for multiple comparisons (resulting in significance of p<0.0083), within the BPD group only there was a negative correlation between plasma AVP and number of errors in recognising threat-related emotions, such that higher AVP corresponded with enhanced responsiveness to negative emotions (average of fear and anger, r=−0.87, p<0.0001, Figure 4d). No correlations were found between copeptin and measures of emotion recognition.

**Figure 4.**
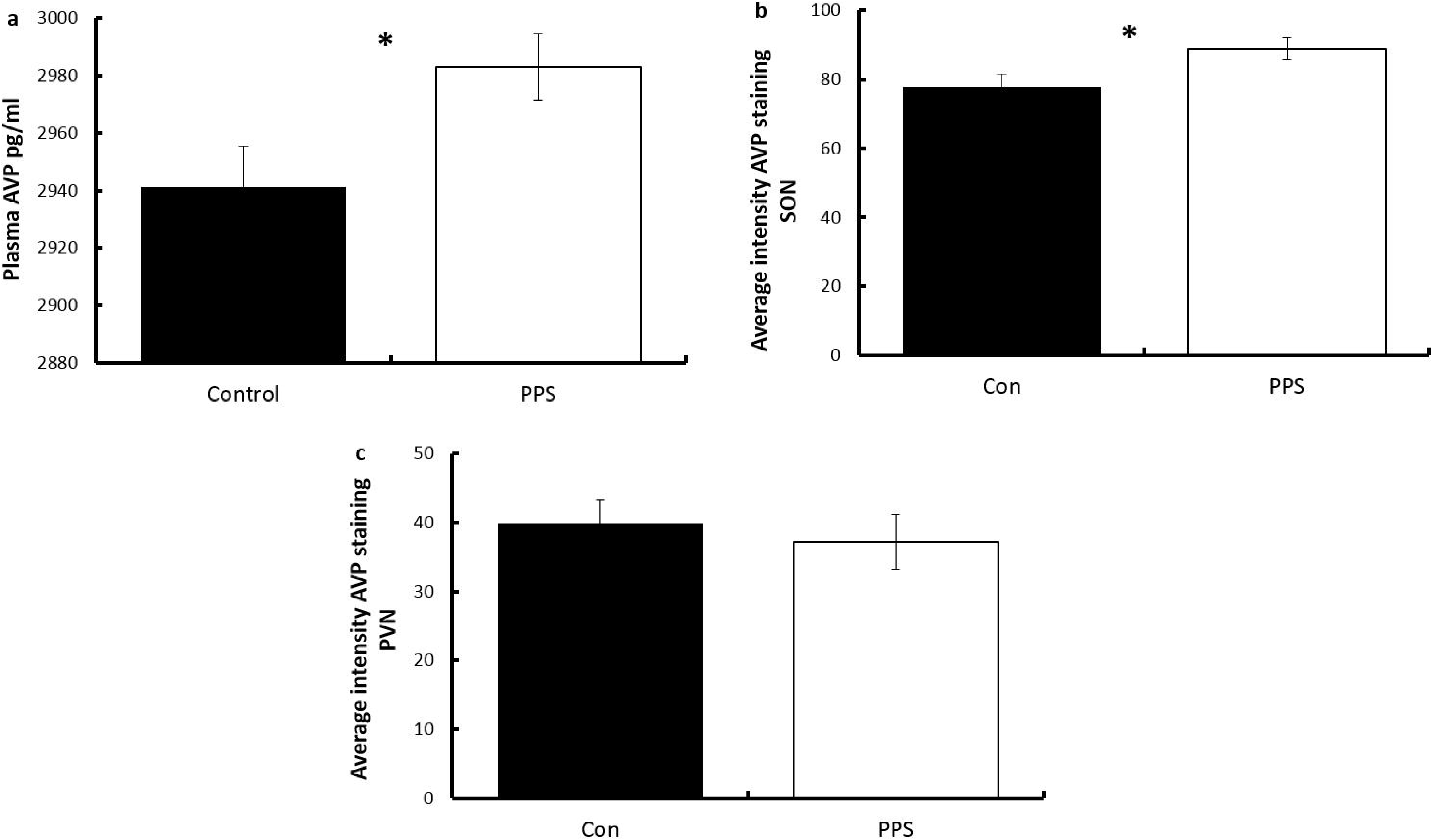
Social behaviour and AVP levels in BPD and controls. BPD participants were a) worse at recognising emotion in the EKMAN task, b) had higher plasma levels of AVP and c) copeptin, and d) within the BPD group only, higher AVP correlated with enhanced detection of negative emotions (fear and anger). *p<0.05. Error bars represent 1 S.E.

## Discussion

Pre-pubertal stress (PPS) resulted in altered social behaviour and elevated protein levels of AVP peripherally in plasma and centrally in the supraoptic nucleus in rats. Relcovaptan, an AVPR1a antagonist, reversed behavioural changes induced by PPS. Similarly, individuals with BPD, a population with a high incidence of childhood adversity, had elevated plasma AVP and altered performance on tests of social emotion recognition. Furthermore AVP levels correlated with increased sensitivity to negative emotional expressions in the BPD group. This suggests the AVP system is at least one mechanism through which early life stress can result in social difficulties in adulthood.

In our animal model, PPS resulted in a sustained increase in AVP levels both peripherally (blood plasma) and centrally (supraoptic nucleus, SON). AVP is mainly synthesised in the magnocellular neurons of the SON and paraventricular nucleus (PVN) and parvocellular neurons of the PVN^63^. Parvocellular neurons project to the zona externa of the median eminence, and AVP originating from these neurons is suggested to be intimately involved in neuro-endocrine HPA axis function at rest and following acute and chronic stress^64, 65^. In contrast, magnocellular neurons project through the zona interna of the median eminence to the posterior pituitary, where AVP is stored in axon terminals and upon stimulation, secreted into systemic blood circulation. This suggests that increased peripheral AVP in our PPS animals is a direct result of the observed upregulation in the magnocellular neurons of the SON. Previously, early life stress in the form of maternal separation increased AVP expression in the PVN of adult mice, but levels in the SON were unaffected^46^. It is unknown whether elevated PVN AVP would produce similar peripheral increases following early life stress as the previous maternal separation study did not examine peripheral AVP levels.

The differential effect of maternal separation and later juvenile stress on AVP in different nuclei of the hypothalamus could result from differences in the stress protocol (short-term physical stress *vs*. maternal separation) or the developmental time point (PND 25-27 *vs*. PND 1-10). This highlights the importance of timing – it is likely that different structures and processes are vulnerable to stressful perturbations at different time points during development. The nature of the stress is also likely to produce different effects in SON and PVN – in adult rats exposure to cold and heat stress increases AVP expression in the PVN, whereas SON AVP only responds to cold stress^66^. Furthermore, differences in the AVP system following early life stress may not be apparent until the system is challenged in a specific manner. AVP was increased in the SON and PVN of maternally separated adult rats and the SON of female prairie voles subjected to chronic social isolation only after experience of stressful intra-sexual social, but not non-social, tests^67-69^.

In contrast to AVP, we found no changes in oxytocin (OXT) following PPS. OXT is also produced in the SON and PVN and responds to certain early life stressors. In rodents, prenatal stress reduces OXT in the PVN, whereas maternal separation has no effect (although OXT receptor binding is altered) ^13, 45, 70-72^. In humans, lower levels of OXT have been detected after severe childhood maltreatment in women, and after early life stress before the age of 12 but not during adolescence in men^73, 74^. Conversely, less severe forms of physical childhood abuse result in increased OXT in adulthood^75^, again highlighting the likely importance of stressor nature and timing for later outcomes.

During a social interaction test, rats exposed to PPS displayed decreased latency to initial contact and decreased duration of individual contact bouts, and number and duration of positively valanced ultrasonic vocalisations were also reduced. Previous work has found that PPS decreases conspecific social interaction in adult males interacting with both adults (rats)^17, 19^ and juveniles (mice)^16^ (but see Tzanoulinou *et al*. 2014^76^). In animals, AVP is important for social behaviour: disruption of AVP neurons impairs social memory specifically, and AVPR1a-/- and AVPR1b-/- mice demonstrate aberrant social recognition^77-79^. AVP infusion into the olfactory bulb and lateral septum facilitates social recognition memory in rats, and along with AVPR1a, AVP modulates aggression through several brain regions^40, 80-83^. This raised the question of whether increased AVP following PPS in our animal model was directly responsible for altered social behaviour. We therefore investigated the ability of an AVPR1a antagonist, Relcovaptan (SR49059) to rescue behaviour following PPS. Social behaviour resulting from PPS was restored by administration of Relcovaptan, suggesting that AVP mediates some of the effects of PPS on adulthood social behaviour, and that this is a reversible phenomenon.

We next investigated whether AVP levels are also elevated in a psychiatric condition associated with high levels of childhood adversity, namely BPD. As with our animal model, we find that AVP protein was elevated in the plasma of BPD participants. Concordant with animal research, AVP and AVPR1a are strongly implicated in human social behaviour. AVPR1a promoter polymorphisms (particularly in RS3 microsatellite repeats) are associated with variation in prosocial behaviour, especially empathy and altruism, and polymorphisms in AVPR1a and AVPR1b have been shown to associate with social dysfunction in autism spectrum disorder ^72, 84-87^. Furthermore, intranasal administration of AVP enhances threat perception and detection of anger in healthy populations, decreases anger detection in schizophrenic men, and improves fear recognition in schizophrenic women ^88-90^. In the present task study, BPD participants demonstrated altered social behaviour. Specifically, they displayed decreased sensitivity to recognising facial expressions of emotion in the Ekman task when compared to controls. This supports previous findings of impaired social cognition in BPD ^10, 32, 36, 58, 60^. We also find that higher AVP is correlated with higher sensitivity to detect fear and anger in this group, suggesting a role for AVP in the recognition of emotions directly related to threat perception in BPD. Interestingly, BPD is a group already known to display heightened attention to threat related social cues^91^. Combined with the results from our animal model, these findings suggest that AVP has a significant role in modulating socio-emotional responses in groups exposed to high levels of childhood trauma.

This translational study provides evidence from across species for the importance of AVP in relation to early life stress and social behaviour. Our results demonstrate that the AVP system may be a useful target for ameliorating social difficulties in conditions associated with high rates of childhood trauma, such as BPD, and that peripheral AVP levels may represent a valuable biomarker for stratification of treatment in such groups.

In conclusion, we find that experience of stressful events early in life is likely to play a profound role in the development of social abnormalities in adulthood, and alterations in the AVP system are at least partially responsible for this effect. AVPR1a is a promising target for intervention in disorders with a social phenotype, especially those produced thorough stressful early life experiences.

## Acknowledgements

We wish to acknowledge support from the Cardiff University Neuroscience and Mental Health Research Institute and The Jane Hodge Foundation who provided NB with fellowship funding during this research, as well as The Waterloo Foundation who provided grant funding for preliminary work (grant number 918-1875). We would also like to thank Dr Oscar Almer for constructing the social testing arena, and Dr Stephane Baudouin for loan of audio recording and analysis equipment. We thank the National Centre for Mental Health (NCMH), which is a collaboration between Cardiff, Swansea and Bangor Universities and is funded by Welsh Government through Health and Care Research Wales. We thank NCMH study participants for their invaluable contribution to this project. The authors declare no competing financial interests.

## References

1. Green MF, Horan WP, Lee J. Social cognition in schizophrenia. Nature Reviews Neuroscience 2015; 16(10): 620–631.

2. Lieb K, Zanarini MC, Schmahl C, Linehan MM, Bohus M. Borderline personality disorder. Lancet 2004; 364(9432): 453–461.

3. Bora E, Pantelis C. Social cognition in schizophrenia in comparison to bipolar disorder: A meta-analysis. Schizophrenia Research 2016; 175(1-3): 72–78.

4. Raine A. Antisocial Personality as a Neurodevelopmental Disorder. Annual Review of Clinical Psychology, Vol 14 2018; 14: 259–289.

5. Green MF, Llerena K, Kern RS. The “Right Stuff” Revisited: What Have We Learned About the Determinants of Daily Functioning in Schizophrenia? Schizophrenia Bulletin 2015; 41(4): 781–785.

6. Javed A, Charles A. The Importance of Social Cognition in Improving Functional Outcomes in Schizophrenia. Frontiers in Psychiatry 2018; 9: 14.

7. Vlad M, Raucher-Chene D, Henry A, Kaladjian A. Functional outcome and social cognition in bipolar disorder: Is there a connection? European Psychiatry 2018; 52: 116–125.

8. Stain HJ, Bronnick K, Hegelstad WTV, Joa I, Johannessen JO, Langeveld J, et al. Impact of Interpersonal Trauma on the Social Functioning of Adults With First-Episode Psychosis. Schizophrenia Bulletin 2014; 40(6): 1491–1498.

9. Palmier-Claus J, Berry K, Darrell-Berry H, Emsley R, Parker S, Drake R, et al. Childhood adversity and social functioning in psychosis: Exploring clinical and cognitive mediators. Psychiatry Research 2016; 238: 25–32.

10. Nicol K, Pope M, Sprengelmeyer R, Young AW, Hall J. Social Judgement in Borderline Personality Disorder. Plos One 2013; 8(11). e73440.

11. Nicol K, Pope M, Romaniuk L, Hall J. Childhood trauma, midbrain activation and psychotic symptoms in borderline personality disorder. Translational Psychiatry 2015; 5. e559.

12. Sandi C, Haller J. Stress and the social brain: behavioural effects and neurobiological mechanisms. Nature Reviews Neuroscience 2015; 16(5): 290–304.

13. de Souza MA, Centenaro LA, Menegotto PR, Henriques TP, Bonini J, Achaval M, et al. Prenatal Stress Produces Social Behavior Deficits and Alters the Number of Oxytocin and Vasopressin Neurons in Adult Rats. Neurochemical Research 2013; 38(7): 1479–1489.

14. Veenema AH, Neumann ID. Maternal separation enhances offensive play-fighting, basal corticosterone and hypothalamic vasopressin mRNA expression in juvenile mate rats. Psychoneuroendocrinology 2009; 34(3): 463–467.

15. Toth M, Halasz J, Mikics E, Barsy B, Haller J. Early social deprivation induces disturdbed social communication and violent aggression in adulthood. Behavioral Neuroscience 2008; 122(4): 849–854.

16. Jacobson-Pick S, Audet MC, Nathoo N, Anisman H. Stressor experiences during the juvenile period increase stressor responsivity in adulthood: Transmission of stressor experiences. Behavioural Brain Research 2011; 216(1): 365–374.

17. Toth E, Avital A, Leshem M, Richter-Levin G, Braun K. Neonatal and juvenile stress induces changes in adult social behavior without affecting cognitive function. Behavioural Brain Research 2008; 190(1): 135–139.

18. Ueno H, Suemitsu H, Murakami S, Kitamura N, Wani K, Matsumoto Y, et al. Juvenile stress induces behavioral change and affects perineuronal net formation in juvenile mice. Bmc Neuroscience 2018; 19. 41.

19. MacKay JC, James JS, Cayer C, Kent P, Anisman H, Merali Z. Protracted Effects of Juvenile Stressor Exposure Are Mitigated by Access to Palatable Food. Plos One 2014; 9(5).

20. Marquez C, Poirier GL, Cordero MI, Larsen MH, Groner A, Marquis J, et al. Peripuberty stress leads to abnormal aggression, altered amygdala and orbitofrontal reactivity and increased prefrontal MAOA gene expression. Translational Psychiatry 2013; 3. e216.

21. Cordero MI, Ansermet F, Sandi C. Long-term programming of enhanced aggression by peripuberty stress in female rats. Psychoneuroendocrinology 2013; 38(11): 2758–2769.

22. Kilian S, Asmal L, Chiliza B, Olivier MR, Phahladira L, Scheffler F, et al. Childhood adversity and cognitive function in schizophrenia spectrum disorders and healthy controls: evidence for an association between neglect and social cognition. Psychological Medicine 2018; 48(13): 2186–2193.

23. Chu DA, Bryant RA, Gatt JM, Harris AWF. Failure to differentiate between threat-related and positive emotion cues in healthy adults with childhood interpersonal or adult trauma. Journal of Psychiatric Research 2016; 78: 31–41.

24. Fernandes V, Osorio FL. Are there associations between early emotional trauma and anxiety disorders? Evidence from a systematic literature review and meta-analysis. European Psychiatry 2015; 30(6): 756–764.

25. Haller J, Harold G, Sandi C, Neumann ID. Effects of Adverse Early-Life Events on Aggression and Anti-Social Behaviours in Animals and Humans. Journal of Neuroendocrinology 2014; 26(10): 724–738.

26. McGrath JJ, McLaughlin KA, Saha S, Aguilar-Gaxiola S, Al-Hamzawi A, Alonso J, et al. The association between childhood adversities and subsequent first onset of psychotic experiences: a cross-national analysis of 23 998 respondents from 17 countries. Psychological Medicine 2017; 47(7): 1230–1245.

27. Kessler RC, McLaughlin KA, Green JG, Gruber MJ, Sampson NA, Zaslavsky AM, et al. Childhood adversities and adult psychopathology in the WHO World Mental Health Surveys. British Journal of Psychiatry 2010; 197(5): 378–385.

28. Beutel ME, Tibubos AN, Klein EM, Schmutzer G, Reiner I, Kocalevent RD, et al. Childhood adversities and distress – The role of resilience in a representative sample. Plos One 2017; 12(3).

29. Johnson JG, Cohen P, Brown J, Smailes EM, Bernstein DP. Childhood maltreatment increases risk for personality disorders during early adulthood. Archives of General Psychiatry 1999; 56(7): 600–606.

30. Ball JS, Links PS. Borderline Personality Disorder and Childhood Trauma: Evidence for a Causal Relationship. Current Psychiatry Reports 2009; 11(1): 63–68.

31. Bailey T, Alvarez-Jimenez M, Garcia-Sanchez AM, Hulbert C, Barlow E, Bendall S. Childhood Trauma Is Associated With Severity of Hallucinations and Delusions in Psychotic Disorders: A Systematic Review and Meta-Analysis. Schizophrenia Bulletin 2018; 44(5): 1111–1122.

32. Preissler S, Dziobek I, Ritter K, Heekeren HR, Roepke S. Social cognition in borderline personality disorder: evidence for disturbed recognition of the emotions, thoughts, and intentions of others. Frontiers in Behavioral Neuroscience 2010; 4.182.

33. Paris J. The etiology of borderline personality disorder – a biopsychosocial approach. Psychiatry-Interpersonal and Biological Processes 1994; 57(4): 316–325.

34. Battle CL, Shea MT, Johnson DM, Yen S, Zlotnick C, Zanarini MC, et al. Childhood maltreatment associated with adult personality disorders: Findings from the collaborative longitudinal personality disorders study. Journal of Personality Disorders 2004; 18(2): 193–211.

35. Wilson S, Stroud CB, Durbin CE. Interpersonal Dysfunction in Personality Disorders: A Meta-Analytic Review. Psychological Bulletin 2017; 143(7): 677–734.

36. Lazarus SA, Cheavens JS, Festa F, Rosenthal MZ. Interpersonal functioning in borderline personality disorder: A systematic review of behavioral and laboratory-based assessments. Clinical Psychology Review 2014; 34(3): 193–205.

37. Albers HE. The regulation of social recognition, social communication and aggression: Vasopressin in the social behavior neural network. Hormones and Behavior 2012; 61(3): 283–292.

38. Albers HE. Species, sex and individual differences in the vasotocin/vasopressin system: Relationship to neurochemical signaling in the social behavior neural network. Frontiers in Neuroendocrinology 2015; 36: 49–71.

39. Caldwell HK, Albers HE. Oxytocin, Vasopressin, and the Motivational Forces that Drive Social Behaviors. Current topics in behavioral neurosciences 2016; 27: 51–103.

40. Caldwell HK. Oxytocin and Vasopressin: Powerful Regulators of Social Behavior. Neuroscientist 2017; 23(5): 517–528.

41. Insel TR. The Challenge of Translation in Social Neuroscience: A Review of Oxytocin, Vasopressin, and Affiliative Behavior. Neuron 2010; 65(6): 768–779.

42. Donaldson ZR, Young LJ. Oxytocin, vasopressin, and the neurogenetics of sociality (vol 322, pg 900, 2008). Science 2009; 323(5920): 1429–1429.

43. Grace SA, Rossell SL, Heinrichs M, Kordsachia C, Labuschagne I. Oxytocin and brain activity in humans: A systematic review and coordinate-based meta-analysis of functional MRI studies. Psychoneuroendocrinology 2018; 96: 6–24.

44. Lefevre A, Hurlemann R, Grinevich V. Imaging neuropeptide effects on human brain function. Cell Tissue Research 2018; 375(1):279–286.

45. Lukas M, Bredewold R, Neumann ID, Veenema AH. Maternal separation interferes with developmental changes in brain vasopressin and oxytocin receptor binding in male rats. Neuropharmacology 2010; 58(1): 78–87.

46. Murgatroyd C, Patchev AV, Wu Y, Micale V, Bockmuhl Y, Fischer D, et al. Dynamic DNA methylation programs persistent adverse effects of early-life stress. Nature Neuroscience 2009; 12(12): 1559–U1108.

47. Veenema AH, Bredewold R, Neumann ID. Opposite effects of maternal separation on intermale and maternal aggression in C57BL/6 mice: Link to hypothalamic vasopressin and oxytocin immunoreactivity. Psychoneuroendocrinology 2007; 32(5): 437–450.

48. Donadon MF, Martin-Santos R, Osorio FD. The Associations Between Oxytocin and Trauma in Humans: A Systematic Review. Frontiers in Pharmacology 2018; 9: 16.

49. Brydges NM, Wood ER, Holmes MC, Hall J. Prepubertal stress and hippocampal function: Sex-specific effects. Hippocampus 2014; 24(6): 684–692.

50. Brydges NM, Whalley HC, Jansen MA, Merrifield GD, Wood ER, Lawrie SM, et al. Imaging Conditioned Fear Circuitry Using Awake Rodent fMRI. Plos One 2013; 8(1). e54197.

51. Brydges NM, Hall L, Nicolson R, Holmes MC, Hall J. The Effects of Juvenile Stress on Anxiety, Cognitive Bias and Decision Making in Adulthood: A Rat Model. Plos One 2012; 7(10). e48143.

52. Jacobson-Pick S, Richter-Levin G. Differential impact of juvenile stress and corticosterone in juvenility and in adulthood, in male and female rats. Behavioural Brain Research 2010; 214(2): 268–276.

53. Hicks C, Ramos L, Reekie T, Misagh GH, Narlawar R, Kassiou M, et al. Body temperature and cardiac changes induced by peripherally administered oxytocin, vasopressin and the non-peptide oxytocin receptor agonist WAY 267,464: a biotelemetry study in rats. British Journal of Pharmacology 2014; 171(11): 2868–2887.

54. Ramos L, Hicks C, Kevin R, Caminer A, Narlawar R, Kassiou M, et al. Acute Prosocial Effects of Oxytocin and Vasopressin When Given Alone or in Combination with 3,4-Methylenedioxymethamphetamine in Rats: Involvement of the V1(A) Receptor. Neuropsychopharmacology 2013; 38(11): 2249–2259.

55. Simola N, Brudzynski SM. Rat 50-kHz ultrasonic vocalizations as a tool in studying neurochemical mechanisms that regulate positive emotional states: Journal of Neuroscience Methods 2018; 310 (1): 33–44.

56. Leichsenring F, Leibing E, Kruse J, New AS, Leweke F. Borderline personality disorder. Lancet 2011; 377(9759): 74–84.

57. Ekman P, Friesen W (1976). Pictures of facial affect. Consulting Psychologists Press, Palo Alto, CA.

58. Unoka Z, Fogd D, Fuzy M, Csukly G. Misreading the facial signs: Specific impairments and error patterns in recognition of facial emotions with negative valence in borderline personality disorder. Psychiatry Research 2011; 189(3): 419–425.

59. Wagner AW, Linehan MM. Facial expression recognition ability among women with borderline personality disorder: Implications for emotion regulation? Journal of Personality Disorders 1999; 13(4): 329–344.

60. Nicol K, Pope M, Hall J. Facial emotion recognition in borderline personality: An association, with childhood experience. Psychiatry Research 2014; 218(1-2): 256–258.

61. Christ-Crain M, Fenske W. Copeptin in the diagnosis of vasopressin-dependent disorders of fluid homeostasis. Nature Reviews Endocrinology 2016; 12(3): 168–176.

62. Balanescu S, Kopp P, Gaskill MB, Morgenthaler NG, Schindler C, Rutishauser J. Correlation of Plasma Copeptin and Vasopressin Concentrations in Hypo-, Iso-, and Hyperosmolar States. Journal of Clinical Endocrinology & Metabolism 2011; 96(4): 1046–1052.

63. Rotondo F, Butz H, Syro LV, Yousef GM, Di Ieva A, Restrepo LM, et al. Arginine vasopressin (AVP): a review of its historical perspectives, current research and multifunctional role in the hypothalamo-hypophysial system. Pituitary 2016; 19(4): 345–355.

64. Aguilera G, Rabadan-Diehl C. Vasopressinergic regulation of the hypothalamic-pituitary-adrenal axis: implications for stress adaptation. Regulatory Peptides 2000; 96(1-2): 23–29.

65. Caldwell HK, Lee HJ, Macbeth AH, Young WS. Vasopressin: Behavioral roles of an “original” neuropeptide. Progress in Neurobiology 2008; 84(1): 1–24.

66. Jasnic N, Dakic T, Bataveljic D, Vujovic P, Lakic I, Jevdjovic T, et al. Distinct vasopressin content in the hypothalamic supraoptic and paraventricular nucleus of rats exposed to low and high ambient temperature. Journal of Thermal Biology 2015; 52: 1–7.

67. Veenema AH, Blume A, Niederle D, Buwalda B, Neumann ID. Effects of early life stress on adult male aggression and hypothalamic vasopressin and serotonin. European Journal of Neuroscience 2006; 24(6): 1711–1720.

68. Perkeybile AM, Bales KL. Early rearing experience is related to altered aggression and vasopressin production following chronic social isolation in the prairie vole. Behavioural Brain Research 2015; 283: 37–46.

69. Perkeybile AM, Bales KL. Early rearing experience is associated with vasopressin immunoreactivity but not reactivity to an acute non-social stressor in the prairie vole. Physiology & Behavior 2015; 147: 149–156.

70. Lee PR, Brady DL, Shapiro RA, Dorsa DM, Koenig JI. Prenatal stress generates deficits in rat social behavior: Reversal by oxytocin. Brain Research 2007; 1156: 152–167.

71. Oreland S, Gustafsson-Ericson L, Nylander I. Short- and long-term consequences of different early environmental conditions on central immunoreactive oxytocin and arginine vasopressin levels in male rats. Neuropeptides 2010; 44(5): 391–398.

72. Avinun R, Israel S, Shalev I, Gritsenko I, Bornstein G, Ebstein RP, et al. AVPR1A Variant Associated with Preschoolers’ Lower Altruistic Behavior. Plos One 2011; 6(9): 5.

73. Heim C, Young LJ, Newport DJ, Mletzko T, Miller AH, Nemeroff CB. Lower CSF oxytocin concentrations in women with a history of childhood abuse. Molecular Psychiatry 2009; 14(10): 954–958.

74. Opacka-Juffry J, Mohiyeddini C. Experience of stress in childhood negatively correlates with plasma oxytocin concentration in adult men. Stress-the International Journal on the Biology of Stress 2012; 15(1): 1–10.

75. Mizuki R, Fujiwara T. Association of oxytocin level and less severe forms of childhood maltreatment history among healthy Japanese adults involved with child care. Frontiers in Behavioral Neuroscience 2015; 9. 138

76. Tzanoulinou S, Riccio O, de Boer MW, Sandi C. Peripubertal stress-induced behavioral changes are associated with altered expression of genes involved in excitation and inhibition in the amygdala. Translational Psychiatry 2014; 4. e410.

77. Bielsky IF, Hu SB, Szegda KL, Westphal H, Young LJ. Profound impairment in social recognition and reduction in anxiety-like behavior in vasopressin V1a receptor knockout mice. Neuropsychopharmacology 2004; 29(3): 483–493.

78. Bielsky IF, Hu SB, Ren XH, Terwilliger EF, Young LJ. The V1a vasopressin receptor is necessary and sufficient for normal social recognition: A gene replacement study. Neuron 2005; 47(4): 503–513.

79. Tobin VA, Hashimoto H, Wacker DW, Takayanagi Y, Langnaese K, Caquineau C, et al. An intrinsic vasopressin system in the olfactory bulb is involved in social recognition. Nature 2010; 464(7287): 413–U110.

80. Dluzen DE, Muraoka S, Landgraf R. Olfactory bulb norepinephrine depletion abolishes vasopressin and oxytocin preservation of social recognition responses in rats. Neuroscience Letters 1998; 254(3): 161–164.

81. Gutzler SJ, Karom M, Erwin WD, Albers HE. Arginine-vasopressin and the regulation of aggression in female Syrian hamsters (Mesocricetus auratus). European Journal of Neuroscience 2010; 31(9): 1655–1663.

82. Bester-Meredith JK, Marler CA. Vasopressin and aggression in cross-fostered California mice (Peromyscus californicus) and white-footed mice (Peromyscus leucopus). Hormones and Behavior 2001; 40(1): 51–64.

83. Delville W, Mansour KM, Ferris CF. Testosterone facilitates aggression by modulating vasopressin receptors in the hypothalamus. Physiology & Behavior 1996; 60(1): 25–29.

84. Knafo A, Israel S, Darvasi A, Bachner-Melman R, Uzefovsky F, Cohen L, et al. Individual differences in allocation of funds in the dictator game associated with length of the arginine vasopressin 1a receptor RS3 promoter region and correlation between RS3 length and hippocampal mRNA. Genes Brain and Behavior 2008; 7(3): 266–275.

85. Uzefovsky F, Shalev I, Israel S, Edelman S, Raz Y, Mankuta D, et al. Oxytocin receptor and vasopressin receptor 1a genes are respectively associated with emotional and cognitive empathy. Hormones and Behavior 2015; 67: 60–65.

86. Cataldo I, Azhari A, Esposito G. A Review of Oxytocin and Arginine-Vasopressin Receptors and Their Modulation of Autism Spectrum Disorder. Frontiers in Molecular Neuroscience 2018; 11: 20.

87. Kim SJ, Young LJ, Gonen D, Veenstra-VanderWeele J, Courchesne R, Courchesne E, et al. Transmission disequilibrium testing of arginine vasopressin receptor 1A (AVPR1A) polymorphisms in autism. Molecular Psychiatry 2002; 7(5): 503–507.

88. Vadas L, Bloch B, Levin R, Shalev I, Israel S, Uzefovsky F, et al. Sex-specific effect of intranasal vasopressin, but not oxytocin, on emotional recognition and perception in schizophrenia patients. European Psychiatry 2017; 41: S387–S388.

89. Guastella AJ, Kenyon AR, Alvares GA, Carson DS, Hickie IB. Intranasal Arginine Vasopressin Enhances the Encoding of Happy and Angry Faces in Humans. Biological Psychiatry 2010; 67(12): 1220–1222.

90. Thompson R, Gupta S, Miller K, Mills S, Orr S. The effects of vasopressin on human facial responses related to social communication. Psychoneuroendocrinology 2004; 29(1): 35–48.

91. Gunderson JG, Herpertz SC, Skodol AE, Torgersen S, Zanarini MC (2018). Borderline personality disorder: Nature Reviews Disease Primers.

